# Matreex: compact and interactive visualisation for scalable studies of large gene families

**DOI:** 10.1101/2023.02.18.529053

**Authors:** Victor Rossier, Clement Train, Yannis Nevers, Marc Robinson-Rechavi, Christophe Dessimoz

## Abstract

Studying gene family evolution strongly benefits from insightful visualisations. However, the evergrowing number of sequenced genomes is leading to increasingly larger gene families, which challenges existing gene tree visualisations. Indeed, most of them present users with a dilemma: display complete but intractable gene trees, or collapse subtrees, thereby hiding their children’s information. Here, we introduce Matreex, a new dynamic tool to scale-up the visualisation of gene families. Matreex’s key idea is to use “phylogenetic” profiles, which are dense representations of gene repertoires, to minimise the information loss when collapsing subtrees. We illustrate Matreex usefulness with three biological applications. First, we demonstrate on the MutS family the power of combining gene trees and phylogenetic profiles to delve into precise evolutionary analyses of large multi-copy gene families. Secondly, by displaying 22 intraflagellar transport gene families across 622 species cumulating 5’500 representatives, we show how Matreex can be used to automate large-scale analyses of gene presence-absence. Notably, we report for the first time the complete loss of intraflagellar transport in the myxozoan *Thelohanellus kitauei*. Finally, using the textbook example of visual opsins, we show Matreex’s potential to create easily interpretable figures for teaching and outreach. Matreex is available from the Python Package Index (pip install matreex) with the source code and documentation available at https://github.com/DessimozLab/matreex.

## Introduction

Studying the evolutionary dynamics of gene families strongly benefits from appropriate visualisation tools. For example, we can draw evolutionary and functional hypotheses by visually correlating gene repertoires with adaptations or between families. Moreover, visualising the evolutionary history of a gene family provides the framework to generalise classical pairwise gene relationships (*e*.*g*. orthology and paralogy) to multiple species (Dunn and Munro 2016). However, the growing number of genomes sequenced and processed by comparative genomic pipelines results in increasingly larger gene families. For example, the OMA database provides families with more than 100’000 members across more than 2’500 species (Altenhoff et al. 2021). Thus, gene family visualisation tools able to integrate this large volume of data and exploit its full potential are needed. Although many tools can represent large gene trees (Xu et al. 2021; Penel and de Vienne 2022), few are interactive, which is essential for users to explore large gene families.

Gene trees labelled with duplications and speciations are typically used to depict the evolutionary history of gene families. However, existing interactive gene tree viewers are not equipped to provide overviews of evolutionary trajectories required to study large gene families spanning thousands of taxa and dozens of subfamilies. To keep gene trees interpretable, most viewers merely rely on collapsing or trimming subtrees, by letting users dynamically expand the relevant ones, while collapsing others (Herrero et al. 2016; Mi et al. 2017; Nguyen et al. 2018; Fuentes et al. 2021). For example, the GeneView of Ensembl collapses by default all subtrees lying outside the lineage of the query gene and provides the option to collapse all nodes at a given taxonomic rank (Herrero et al. 2016). Similarly, the PhyloView of Genomicus displays the gene tree at a user-defined taxon and provides many customization features such as trimming outgroups (relative to the query gene) or duplication nodes (Nguyen et al. 2018). However, a collapsed or trimmed subtree is mostly uninformative, as its gene content and topology is not shown. Therefore, users can only choose between keeping a complete and often intractable gene tree, or collapsing nodes and hiding the information of its children, with no middle ground. Moreover, these viewers are limited by their slow reactivity, which makes the exploration of large gene trees cumbersome. For example, a couple of seconds is needed to collapse a node in Ensembl GeneView or PhylomeDB, while any action brings the user back to the top of the page in Genomicus PhyloView. Faster and more scalable web-based tools have been introduced to visualise large phylogenies of species or of viral genomes (Robinson et al. 2016; Turakhia et al. 2020), but they are not tailored to display gene families and also lack a way to summarise relevant information contained in the different relevant parts of the phylogenies.

Alternatively, gene families can be represented as vectors of gene copy numbers across species or phylogenetic profiles. Although these were initially developed to infer gene functions, as repeated cooccurrences provide evidence of interaction (Pellegrini et al. 1999), visualising these profiles has proven useful to illustrate the gene content of extant species (Musilova et al. 2021; Horn et al. 2022) or to compare likely coevolving families (van Dam et al. 2013; Nevers et al. 2017). Indeed, displaying the full gene repertoire of a species in the same column (or row) and all gene family members in the same row (or column) enables rapid visual identification of repeated and correlated gene presence and absences. The relevance of this kind of compact representation of gene families is evidenced by the large number of tools developed for that task (Sadreyev et al. 2015; Cromar et al. 2016; Tran et al. 2018; Tremblay et al. 2021; Ilnitskiy et al. 2022). However, unlike gene trees, phylogenetic profiles do not show evolutionary relationships among the genes; for instance, it is not possible to deduct from a profile alone whether two gene absences are the result of independent losses, or a single loss in a common ancestor.

Here, we introduce Matreex, an innovative viewer for large gene families that bridges the gap between these two typical representations of gene families: gene trees that provide their complete evolutionary picture but can be cumbersome to read and phylogenetic profiles that efficiently depict the distribution of genes across species but lack the evolutionary component. Matreex builds on the reactive framework from the Phylo.IO viewer (Robinson et al. 2016) and integrates phylogenetic profiles to summarise collapsed subtrees. Thus, it simplifies gene tree visualisation while reducing the information loss. The resulting highly compact and reactive visualisation of evolution enables Matreex to scale-up to the ongoing deluge of genomic data. Moreover, it provides the opportunity for new biological discoveries, for the production of paper figures, and for didactic support for teaching in evolutionary biology. We illustrate Matreex with three biological applications.

## New Approach

To enable compact and reactive visualisation of large gene families, Matreex complements the gene tree with a matrix of phylogenetic profiles and a species tree (Fig. 1). Thus, when collapsing a subtree to simplify the gene tree, the distribution of gene copy numbers across species remains available in the corresponding row. This keeps information about ancestral events such as gene loss, duplication or transfer. Moreover, the species tree displayed orthogonally from the gene tree provides what is often a good proxy for the topology of these subtrees (Morel et al. 2020). Remarkably, the extreme case, where all subtrees without duplications and congruent with the species tree are collapsed, provides the same information as a fully extended gene tree but much more compact. Indeed, these subtrees do not need to be displayed because their topology is explicit in the species tree We provide rapid access to this view with Matreex’s “Smart Collapse” option.

**Figure 1.**
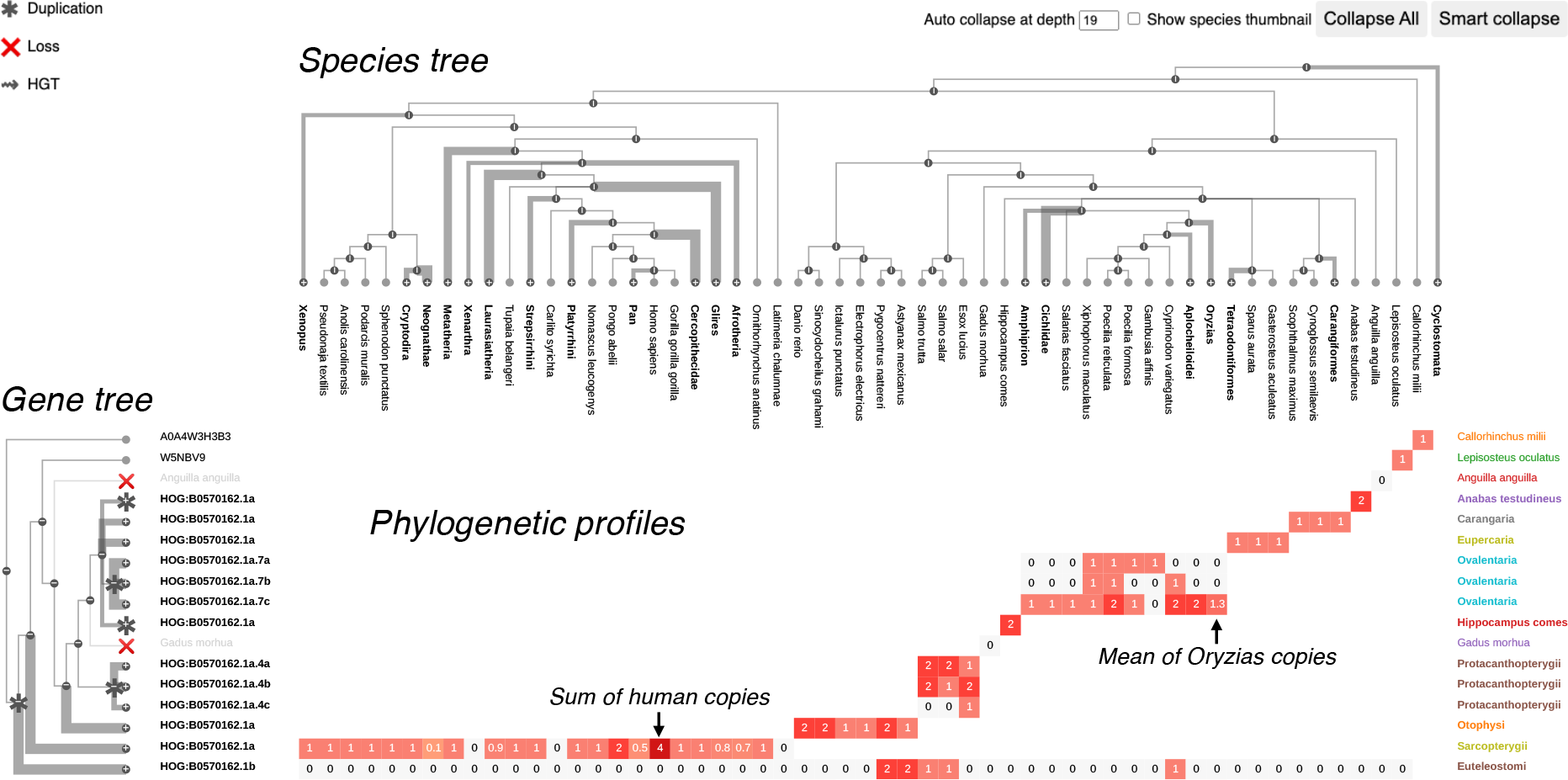
Matreex’s layout consists of a gene tree, a species tree and a matrix of phylogenetic profiles. Gene tree labels represent gene subfamily memberships (OMA HOGs in this case) for collapsed nodes, gene ids for leaves, and taxon or species names for lost genes (implied from the species tree). Branch thickness increases with the collapsed subtree size and cell colour darkness with the number of in-paralogs in the cell. The taxonomic levels of collapsed subtrees and phylogenetic profiles are annotated on the right. “Auto-collapse at depth” enables the automatic collapse of the species tree at a given depth from the root. “Show species thumbnail” enables displaying a taxon image (at present from Wikipedia) when hovering over a taxon. “Collapse All” and “Smart collapse” are two default views described in the main text. The examples shown are red-sensitive visual opsins (data from OMA, All.Dec2021 release). Italic annotations do not belong to the Matreex layout but were added for figure clarity.

For genes families with a high number of duplication events, collapsing only subtrees without duplication is not enough and summarising them requires also collapsing subtrees with subfamilies (children of duplication nodes). In that case, the resulting phylogenetic profiles depict the combined gene content of each subfamily per species. For example, the profile of the collapsed *Sarcopterygii* subtree of the red-sensitive visual opsin (LWS) family shows four copies for humans due to the existence of primates-specific subfamilies (Fig. 1). To deal with large gene families, Matreex includes the option of collapsing all subfamilies, including the root node (Matreex’s “Collapse All”), as manually collapsing many nodes can be tedious. Starting from the family phylogenetic profile, the user can then unfold more and more specific subfamilies, thus revealing their species distributions and gene copy number variations. In particular, unfolding a node will reveal the gene tree topology until the next duplication nodes, which define the child subfamilies. Other subtrees will remain collapsed, as their topology is redundant with the species tree. This approach is user-friendly because it begins with a highly summarised view of the family before zooming into more specific subfamilies of interest.

Two main processes increase the size of gene families in practice: gene duplications and the increase in the number of species. The latter increases both with the number of available genomes and with the progress of orthology assessment methods and resources in handling a growing number of species (*e*.*g*. (Kriventseva et al. 2019; Altenhoff et al. 2021; Cantalapiedra et al. 2021; Rossier et al. 2021)). Thus, the ability to control which species (or taxa) to show and which to hide is key to allow users to zoom on taxa of interest, while achieving high levels of gene family compactness. For that task, Matreex provides control over which taxa are displayed through its interactive species tree. When collapsing a taxon in the species tree, all corresponding gene tree nodes are also collapsed, and the phylogenetic profiles are summarised. This is done by averaging the numbers of in-paralogs of the species descending from the collapsed node. For example, collapsing the *Oryzias* node in the LWS family automatically merges *Oryzias*-specific subfamilies and averages the copy numbers of *Oryzias* species (Fig. 1). Moreover, to facilitate the exploration of large species trees, Matreex provides the option to collapse every taxon after a given node depth from the root.

Finally, Matreex implements several other design features to further facilitate the user experience. First, as scientific names can be quite obscure, images are displayed when hovering over taxon labels; at present Wikipedia images are used but other sources could be easily implemented. Second, to highlight lineage-specific expansions, the matrix of phylogenetic profiles is displayed as a heatmap for which custom colours can be used to highlight specific clades.

## Availability and implementation

Matreex available from the Python Package Index (pip install matreex) with the source code, the documentation and code to reproduce the below figures available at https://github.com/DessimozLab/matreex.

Matreex is both a command-line tool and a python library that produces html output files, which can be viewed in standard web browsers and therefore easily shared. It is implemented in JavaScript with the D3 library and wrapped in a Python module that supports the OMA and PANTHER APIs (Kaleb et al. 2019; Mi et al. 2021). For these two databases, only the gene family identifier, or list of gene family identifiers, is required as input. Moreover, users can also upload their own gene tree in JSON (format described in the GitHub). However, similarly to OMA HOGs or PANTHER gene trees, Matreex requires input gene trees to be consistent with their associated species trees (where speciations follow the same order in both trees). To obtain a tree fulfilling this condition, we suggest using a gene tree inference method that already attempts to minimise conflicts between gene trees and species trees (Boussau et al. 2013; Noutahi et al. 2016; Morel et al. 2020), and to replace remaining conflicting regions with polytomies.

## Applications

In this section, we illustrate how Matreex facilitates the analysis of gene families on three different use-cases with real biological applications.

### Origin and evolution of eukaryotic MutS genes

Matreex enables users to analyse precisely the gene repertoire evolution of large multi-copy families. First, subfamily gene repertoires can be correlated among themselves or with adaptations. Secondly, the gene tree enables to study the evolutionary relationships between phylogenetic profiles. This can be useful, for instance, to differentiate orthologous from paralogous profiles and to diagnose the underlying gene tree. In this last application, we performed a detailed analysis of the MutS family, whose evolutionary history remains largely under debate. Specifically, we used Matreex to simplify the task of systematically contrasting existing knowledge with the data at hand (Fig. 2). First, we evaluated whether some established hypotheses were further evidenced or challenged by the examined MutS gene tree. Second, we assessed which still-debated hypotheses were supported by this tree and, third, we formulated new hypotheses by searching for patterns in this new visualisation. Finally, we contextualised the results with functional and evolutionary knowledge from the literature to highlight the importance of such an approach.

**Figure 2.**
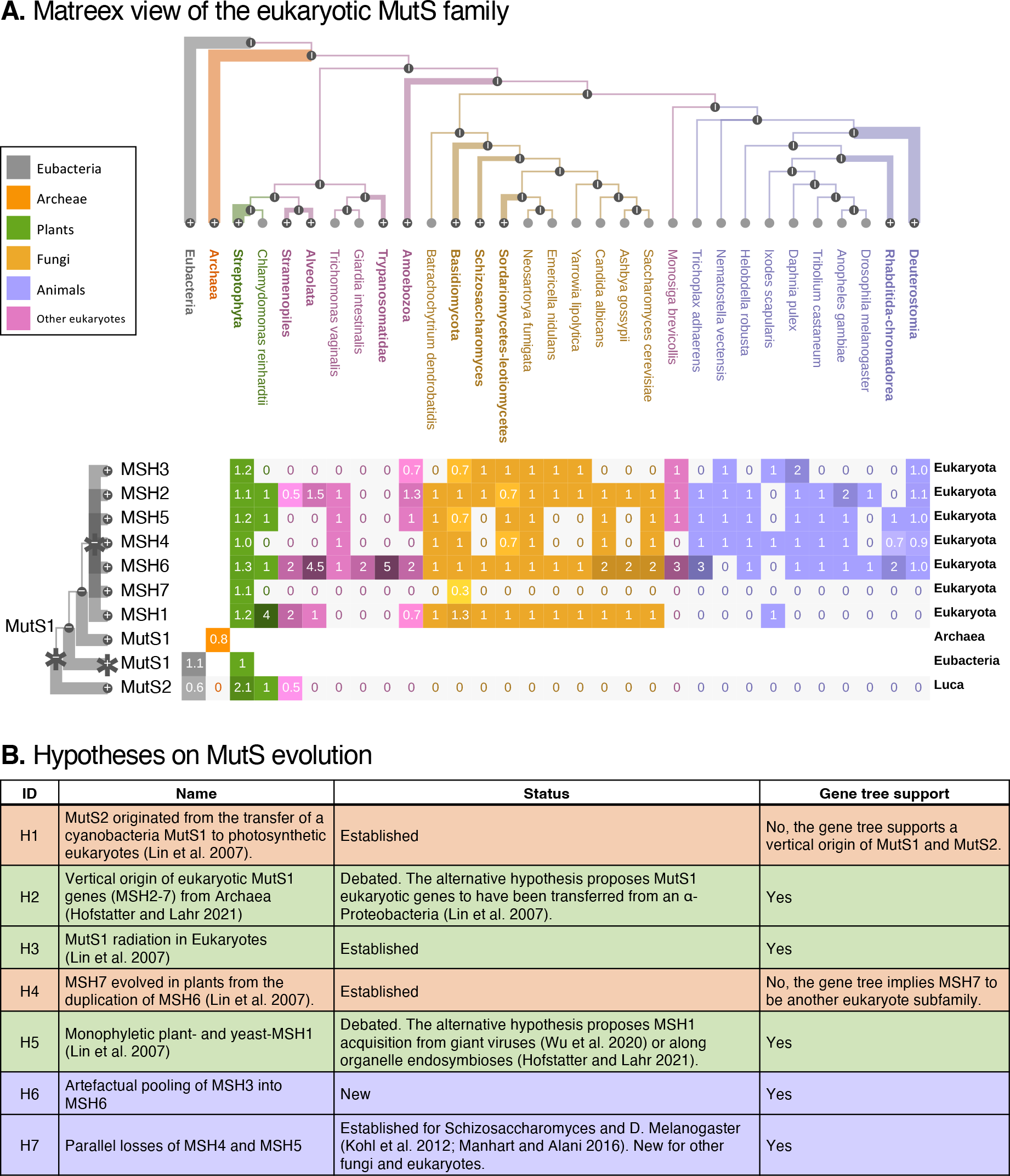
Detailed evolutionary analysis of the eukaryotic MutS family. A. Matreex view of the MutS gene family (gene tree from PANTHER v.17). Clade legends and gene family names do not belong to the Matreex layout but were added for figure clarity. B. Hypotheses on MutS evolution discussed in the text with their level of support in the literature and in the examined gene tree. An orange, green, or blue background indicates, respectively, a conflict with the literature, no conflict with the literature, or a new hypothesis from this work.

MutS genes are involved in the DNA mismatch repair pathway (Liu et al. 2017; Mi et al. 2021). Although bacteria have multiple MutS genes, only MutS1 and MutS2 are found in both bacteria and eukaryotes. MutS2 is found only in photosynthetic eukaryotes and its transfer from cyanobacteria through the chloroplast endosymbiosis is well established (Fig. 2, H1) (Lin et al. 2007). Matreex shows clearly the absence of MutS2 in other eukaryotes and *Archaea*, as well as its two copies in plants (*Streptophyta*). However, PANTHER predicts a vertical origin of MutS2 from a pre-LUCA duplication followed by independent losses in *Archaea* and non-photosynthetic eukaryotes. Matreex shows this evolutionary trajectory with a fully connected phylogenetic profile for MutS2 and losses instead of empty cells for *Archaea* and most eukaryotes.

In contrast to MutS2, the origin of MutS1 remains under debate. Eukaryotic MutS1 genes (MSH2-7) were first thought to originate from the mitochondria endosymbiosis of an α*-Proteobacteria* (Lin et al. 2007) until the Asgard *Archaea* MutS1 was found to be more closely related to Eukaryotes than to α*-Proteobacteria* (Hofstatter and Lahr 2021). This implies the vertical origin of MSH2-7 from *Archaea* (Fig. 2, H2). Matreex shows clearly these orthologous relationships between archeal MutS1 and eukaryotic MSH2-7 because they form a monophyletic clade in the gene tree and their profiles do not overlap.

MSH2-6 genes originated from duplications in the eukaryote ancestor (Fig. 2, H3), while MSH7 arose from a plant-specific duplication of MSH6 (Fig. 2, H4) (Lin et al. 2007). Matreex clearly represents this radiation with a compact block of subfamily profiles, although PANTHER supports MSH7 to be another eukaryote subfamily. This is visible with Matreex by its phylogenetic profile mainly displaying zeros instead of empty cells. Given the improbable number of implied lost and the current state of the literature, this pattern most likely reflects a methodological artefact rather than a true evolutionary scenario.

However, the origins of the plant- and fungi-MSH1 remain unclear. Although originally thought to descend from the same MutS ancestor as MSH2-7 (Fig. 2, H5) (Lin et al. 2007), an acquisition of MSH1 in fungi along with the mitochondrial endosymbiosis has been suggested (Hofstatter and Lahr 2021). Similarly, the plant-MSH1 could have been acquired from giant viruses or along the chloroplast endosymbiosis (Wu et al. 2020; Hofstatter and Lahr 2021). The underlying gene tree supports the original hypothesis as we found the plant- and fungi-MSH1 genes in the same eukaryotic subfamily. Matreex helped to draw this conclusion as both plant- and fungi-MSH1 belong to the same collapsed subtree and phylogenetic profile, indicating a monophyletic origin. Moreover, we recovered the absence of MSH1 in animals (with the exception of the tick *Ixodes scapularis* whose copy likely originated from transfer or contamination from the genome of its *Rickettsia* endosymbiont), which has been recently linked with the exceptionally high evolutionary rates of their mitochondrial genes, as MSH1 is involved in repairing their sequences (Wu et al. 2020).

Matreex simplifies the identification of gene repertoire evolutionary patterns. Thus, we observed unexpected expansions of MSH6 in eukaryotes (*e*.*g. Alveolata, Trypanosomatidae*), yeast (*Saccharomyces cerevisiae*), nematodes and fruitfly (*Drosophila Melanogaster*). Then, by expanding the gene tree, we noticed many MSH3 genes misclassified as MSH6, which coincides with predicted MSH3 losses. For example, of the five *Trypanosomatidae* copies, two were surely misclassified MSH3 and MSH5 genes, and one was undefined. Thus, although the loss of MSH3 in nematodes and insects and the trypanosome-specific MSH8 subfamily are documented (Bell et al. 2004; Muthye and Lavrov 2021), we hypothesised that MSH6 is artefactually attracting other genes, in particular MSH3 ones, during phylogenetic reconstruction (Fig. 2, H6).

Finally, we observed repeated and correlated losses of MSH4 and MSH5 in fungi, fruit fly and other eukaryotes (Fig. 2, H7). While losses in the latter are likely artefactual (Rzeszutek et al. 2022), *Schizosaccharomyces* and *D. Melanogaster* are known to have lost and replaced MSH4 and MSH5 for meiotic recombination (Kohl et al. 2012; Manhart and Alani 2016). Moreover, given that these two genes form an obligate complex, other correlated losses in fungi are plausible and could provide good candidates to study alternative meiotic recombination mechanisms.

### Coevolution of the intraflagellar transport genes

Matreex enables to perform gene presence-absence analyses for dozens of non-homologous families spanning hundreds of species in a few minutes. This is useful to visualise the result of a phylogenetic profile search (Altenhoff et al. 2021) or to study coevolving gene families (*e*.*g*. involved in the same pathway). In this second application, we illustrate the latter by generalising a study on eukaryotic intraflagellar transport (IFT) genes from 622 species, compared to the 52 used originally (van Dam et al. 2013). Specifically, we used Matreex to simplify the task of contrasting our results with the literature and to propose new biological hypotheses.

Eukaryotic flagella (cilia) are involved in cell motility and sensory detection (Nevers et al. 2017). Their dysfunction is the cause of ciliopathies in humans (Badano et al. 2006). The IFT complex is essential to build and maintain the flagella. From an evolutionary perspective, IFT is a great example of the “last-in, first-out” hypothesis (van Dam et al. 2013), whereby modules added last are more dispensable and thus, lost first. Indeed, of the three IFT modules (IFT-α, IFT-β and BBSome), BBSome and IFT-α emerged from IFT-β duplications and their loss often precedes the complete loss of IFT and cilia. Thus, studying how ciliated eukaryotes cope with partial IFT loss is promising for the treatment of IFT-related human ciliopathies such as the Bardet–Biedl syndrome caused by BBSome alterations (Badano et al. 2006).

Repeated and correlated losses are visible at a glance in Matreex with columns of zeros on light grey backgrounds (Fig. 3). As expected, complete loss of IFT complexes were detected in the main nonciliated taxa (*e*.*g. Spermatophyta, Dikarya* or *Amoebozoa*). Moreover, due to the sheer number of used genomes, summarised in one easy to read figure, we were able to identify many other complete IFT losses. Although most were already established (*e*.*g. Fonticula alba, Creolimax fragrantissima, Capsaspora owczarzaki* (Torruella et al. 2015), *Entamoeba* (Wickstead and Gull 2007)), we report the first evidence to our knowledge of a complete loss of IFT in the mixozoan *Thelohanellus kitauei*, likely indicating the loss of the organelle in this species.

**Figure 3.**
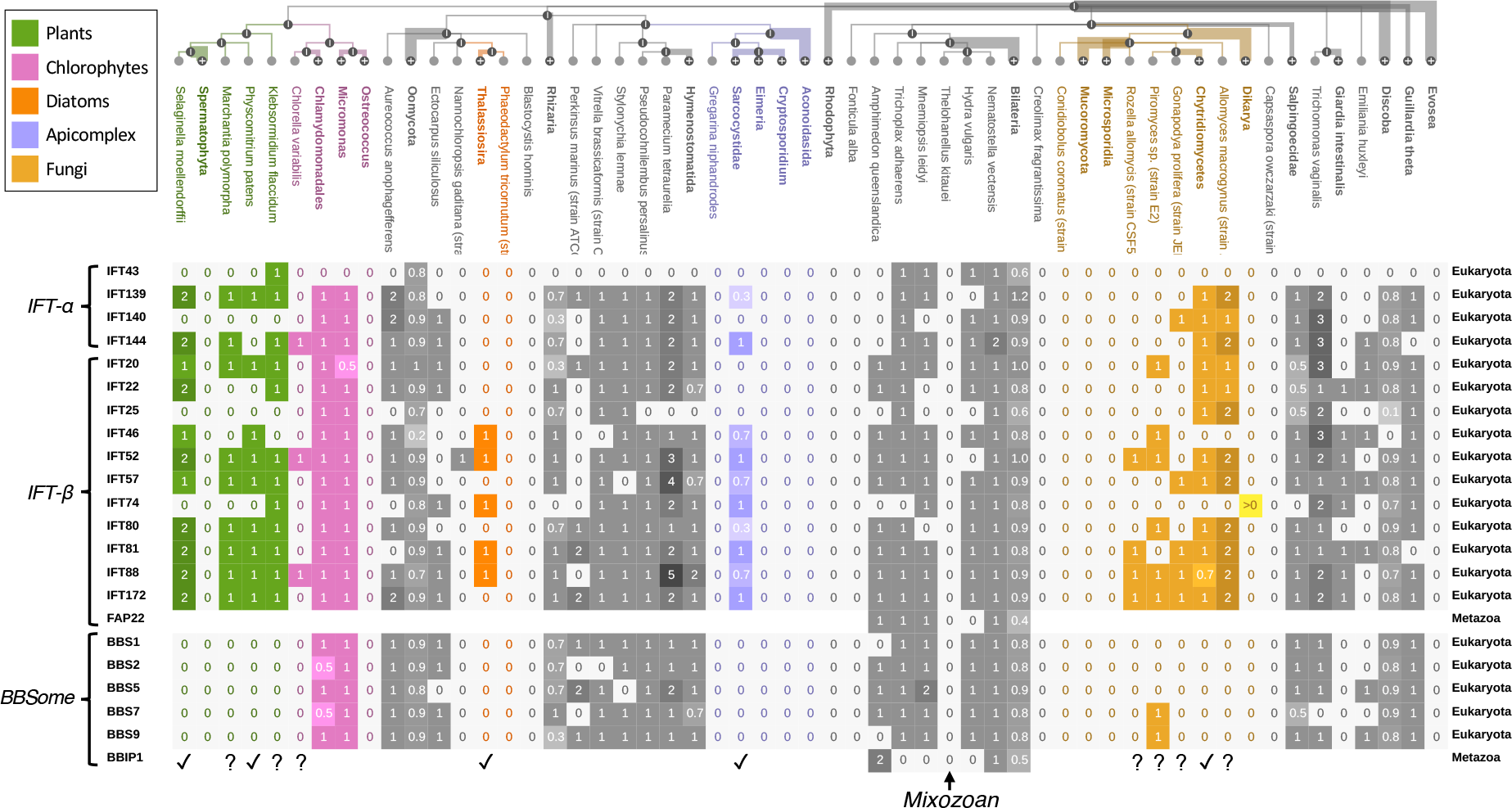
Intraflagellar transport gene families. (data from OMA All.Dec2021). Colored clades display partial and complete IFT losses that fit the “last-in, first-out” hypothesis for gene module evolution. ✓ highlight partial IFT losses reported by van Dam et al. (2013) and ?, the ones reported here. We also report the first evidence to our knowledge of a complete loss of IFT in the mixozoan *Thelohanellus kitauei*. Clade legends, Italic annotations, brackets, ✓ and ? symbols do not belong to the Matreex layout but were added for figure clarity.

Then, we could first quickly confirm all established patterns of BBSome and IFT-α losses in species closely related to non-ciliated clades with complete IFT loss from (van Dam et al. 2013). Specifically, we detected the loss of BBSome in basal plants (*Selaginella moellendorffii* and the moss *Physcomitrella patens*) close to seed plants (*Spermatophyta*), in the apicomplexa *Sarcocystidae* (*Toxoplasma gondii* clade) close to *Aconoidasida* (*Plasmodium falciparum* clade) and in the basal fungi *Chytridiomycetes* (*Batrachochytrium dendrobatidis* clade) close to *Dikarya* and *Mucoromycota*. We also recovered the loss of BBSome and IFT-α in the diatoms *Thalassiosira* close to *Phaeodactylum tricornutum*. Secondly, we could identify other independent losses supporting the “last-in, first-out” hypothesis. In particular, we found two losses of BBSome in the basal plants *Marchantia polymorpha* and *Klebsormidium flaccidum*. We also identified complete IFT losses in another three apicomplexa clades (*Eimeria, Cryptosporidium* and *Gregarina niphandrodes*) and two basal fungi clades (*Microsporidia* and *Conidiobolus coronatus*). Moreover, we found evidence for losses of BBSome and IFT-α in four basal fungi. While *Rozella allomycis* lacks all BBSome and IFT-α genes, *Piromyces sp*. and *Gonapodya prolifera* were found with merely one IFT-α and two BBSome genes, respectively. *Allomyces macrogynus* lacked BBSome. Finally, the presence of one IFT-α and two IFT-β genes in the chlorophytes *Chlorella variabilis* close to *Ostreococcus* provides a new candidate replicate for this “last-in, first-out” hypothesis. Its low number of IFT genes, which indicates dysfunctional cilia, could be due to the endosymbiont nature of *Chlorella variabilis (Blanc et al. 2010)*.

When many gene families underwent duplications in the same species, the column attracts the eye as it becomes darker in Matreex. Thus, we identified four species with many duplicates of IFT-α and IFT-β genes. Although *Paramecium tetraurelia* and *Trichomonas vaginalis* have undergone whole genome duplications (Aury et al. 2006; Carlton et al. 2007), *Paramecium tetraurelia* IFT57 copies show evidence of subfunctionalization (Shi et al. 2018), while *Trichomonas vaginalis* displays specialised cilia that could have required the recruitment of additional IFT copies. Finally, to explain the retention of *S. moellendorffii* and *Allomyces macrogynus* duplicates, that have lost BBSome, we may speculate whether these extra copies could have been co-opted to replace the BBSome functions.

### The visual opsin gene repertoire correlates with adaptations in vertebrates

Matreex enables to quickly identify correlations between adaptations, or phenotypes, and variations in gene copy numbers. Matreex further facilitates the task by depicting losses on a light grey background and expansions on a darker one. Clades can also be coloured to highlight the correlation (using the Matreex library examples available on GitHub). Here, we illustrate this feature with the textbook example of visual opsins (Graur and Li 1999; Paul G. Higgs and Teresa K. Attwood 2005). Because Matreex’s representation facilitates an intuitive interpretation of the data, we expect this usage to become particularly popular in outreach tasks including teaching and conference presentations.

The vertebrate ancestor had one rod opsin (Rhodopsin) for dim light vision and four cone opsins for a tetrachromatic vision, each sensitive to a specific range of light wavelength (Musilova et al. 2021). Specifically, the shortest wavelengths are absorbed by the violet-sensitive opsin (SWS1), followed by the blue- (SWS2) and green-sensitive (RH2) opsins for intermediate wavelengths. The red-sensitive (LWS) opsins absorb for the largest ones. By contrast, mammals and snakes lack the blue- and greensensitive opsins, likely due to the nocturnal lifestyle of their ancestors (Borges et al. 2018; Katti et al. 2019). However, old-world primates (*Catarrhini*, including humans) regained a more complex colour vision by co-opting a red-sensitive opsin duplicate to absorb green wavelengths, which possibly helped primates to identify edible fruits (Carvalho et al. 2017). Matreex shows clearly and at a glance both the losses in mammals and snakes (Fig. 4, pink), as series of light grey background zeroes, and the secondary amplification in *Homo sapiens* and *Pongo abelii* (Fig. 4, green), as darker cells with larger numbers of genes.

**Figure 4.**
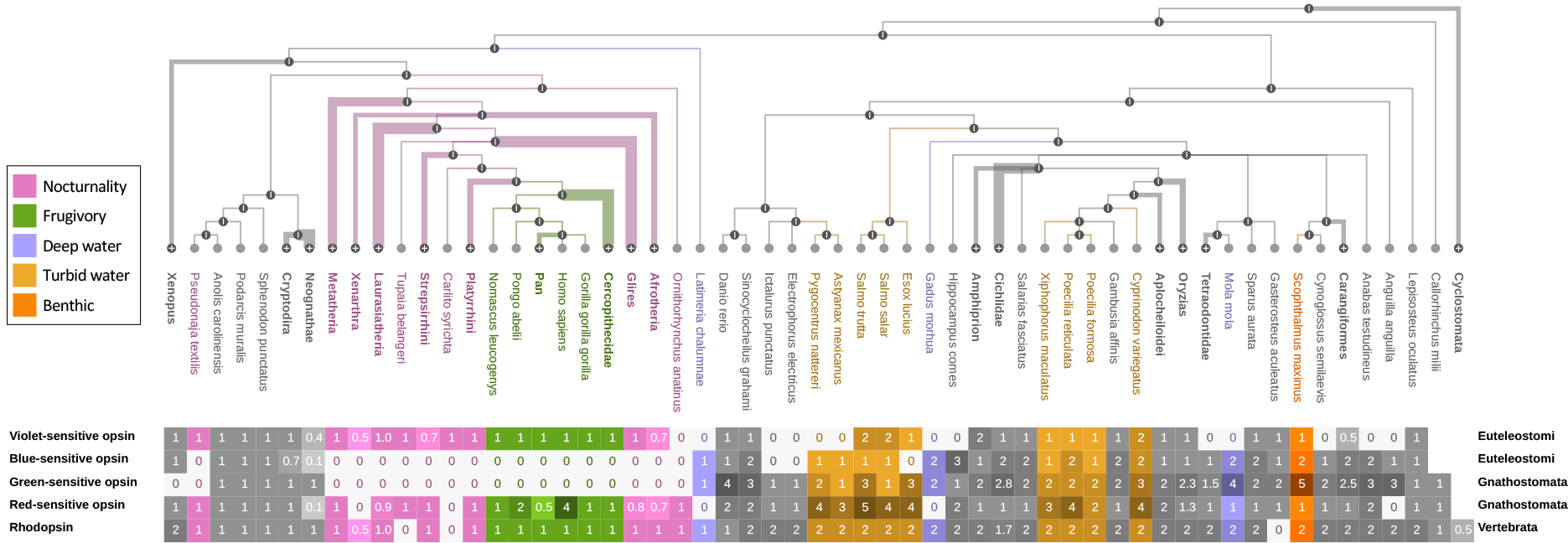
Visual opsin families in vertebrates. (data from OMA All.Dec2021). Adaptations involved in textbook correlations with patterns of gene losses and duplications are annotated with separate colours. Nocturnality in snakes and mammals: loss of blue- and green-sensitive opsins. Frugivory in old-world primates: duplications of red-sensitive opsin. Deep-water: loss of violet- and red-sensitive opsins, duplications of blue- and green-sensitive opsins. Turbid water: duplications of red-sensitive opsins. Benthic: duplications of green-sensitive opsins. Ecological niche legends on the left do not belong to the Matreex layout but were added for figure clarity.

By contrast, the visual opsin repertoire of fishes is much more variable, likely due to the diversity of underwater light environments (Musilova et al. 2021) and this is immediately visible in the Matreex representation. In deep water, the light spectrum is shrinked to absorb only blue and green. Thus, deeper-living species are expected to lose red- and violetsensitive opsins, while duplicating the green- and blue-sensitive ones to compensate for the lower photon abundance. Here, such an evolutionary pattern was detected in the cod (*Gadus morhua*, depth: 150-200m, max. 600m), the sunfish (*Mola mola*, depth 30-70m, max. 480m) and the coelacanth (*Latimeria chalumnae*, depth: 180-250m, max. 700m) (Fig. 4, purple). Moreover, we found the most green-sensitive opsins (five) in the turbot flatfish (*Scophthalmus Maximus*), which could be an adaptation to deep benthic life, as previously suggested (Wang et al. 2021) (Fig. 4, orange). Conversely, the light spectrum is shifted toward longer wavelengths in turbid water, thus favouring red-opsin duplications (Musilova et al. 2021). The present gene tree supports this assumption as we found the most red-opsin copies in fishes that live in the turbid freshwater and brackish habitats (Fig. 4, brown). In particular, five copies were detected in the brown trout (*Salmo trutta*) and four in the red piranha (*Pygocentrus nattereri*), the atlantic salmon (*Salmo salar*), the northern pike (*Esox lucius*), the guppy (*Poecilia reticulata*) and the pupfish (*Cyprinodon variegatus*).

## Conclusion

At a time when the goal of sequencing all eukaryotic species before 2030 has been set (Lewin et al. 2022), it has become critical to develop new methods to represent this huge volume of upcoming data. Here, we introduced an innovative tool to scale the visualisation of gene families and illustrate its usefulness with three biological applications. First, we demonstrated Matreex’s usefulness in delving into precise evolutionary analyses of multi-copy gene families by combining the gene tree with phylogenetic profiles. Secondly, by displaying 22 Intraflagellar gene families across 622 species cumulating 5’500 representatives, we showed how Matreex can be used for analyses of gene presence-absence and reported for the first time the complete loss of IFT in the mixozoan *Thelohanellus kitauei*. Finally, using the textbook example of visual opsins, we demonstrated Matreex’s potential to create easily interpretable figures for outreach tasks. Thus, we hope Matreex will become a valuable tool to gain insights into the evolution of increasingly large gene families.

